# Screening performance of abbreviated versions of the UPSIT smell test

**DOI:** 10.1101/443127

**Authors:** Theresita Joseph, Stephen D. Auger, Luisa Peress, Daniel Rack, Jack Cuzick, Gavin Giovannoni, Andrew Lees, Anette E. Schrag, Alastair J. Noyce

## Abstract

**Background:** Hyposmia features in several neurodegenerative conditions, including Parkinson’s disease (PD). The University of Pennsylvania Smell Identification Test (UPSIT) is a widely used screening tool for detecting hyposmia, but is time-consuming and expensive when used on a large scale.

**Methods:** We assessed shorter subsets of UPSIT items for their ability to detect hyposmia in 891 healthy participants from the PREDICT-PD study. Established shorter tests included Versions A and B of both the 4-item Pocket Smell Test (PST) and 12-item Brief Smell Identification Test (BSIT). Using a data-driven approach, we evaluated screening performances of 23,231,378 combinations of 1-7 smell items from the full UPSIT.

**Results:** PST Versions A and B achieved sensitivity/specificity of 76.8%/64.9% and 86.6%/45.9% respectively, whilst BSIT Versions A and B achieved 83.1%/79.5% and 96.5%/51.8% for detecting hyposmia defined by the longer UPSIT. From the data-driven analysis, two optimised sets of 7 smells surpassed the screening performance of the 12 item BSITs (with validation sensitivity/specificities of 88.2%/85.4% and 100%/53.5%). A set of 4 smells (Menthol, Clove, Gingerbread and Orange) had higher sensitivity for hyposmia than PST-A, -B and even BSIT-A (with validation sensitivity 91.2%). The same 4 smells also featured amongst those most commonly misidentified by 44 individuals with PD compared to 891 PREDICT-PD controls and a screening test using these 4 smells would have identified all hyposmic patients with PD.

**Conclusion:** Using abbreviated smell tests could provide a cost-effective means of screening for hyposmia in large cohorts, allowing more targeted administration of the UPSIT or similar smell tests.

## INTRODUCTION

Reduced ability to detect and recognise smells (hyposmia) commonly occurs with increasing age. When profound it can be a feature of several neurodegenerative disorders, including Parkinson’s disease (PD)(1,2). Hyposmia is observed in up to 90% of PD patients(3), and is considered a sensitive non-motor symptom for discriminating between PD patients and healthy controls(4). The onset of hyposmia can predate motor symptoms by years(5,6), and is associated with an increased risk of being diagnosed with PD(7–9). The neural substrate behind olfactory dysfunction in PD is incompletely understood, however neuropathological evidence points to the olfactory bulb being among the first regions to demonstrate neuronal loss and accumulation of intracytoplasmic a-synuclein rich Lewy bodies(10–12), before the pathology involves more central regions. Early olfactory dysfunction is also implicated in Alzheimer’s disease (AD) and other neurodegenerative diseases(13–15), making it a potential marker for the early identification of these processes(16–18).

Several tests have been created to screen for olfactory dysfunction; assessing the identification, discrimination and detection thresholds for different odours(2,19). The University of Pennsylvania Smell Identification Test (UPSIT), developed by Sensonics, is the most commonly-used smell test worldwide(20), and comprises 40 “scratch-and-sniff” microencapsulated odorant strips divided across 4 booklets (10 in each). For each strip, participants are required to identify the correct smell from a forced choice of 4 possible answers. The total number of smells correctly identified out of 40 is then compared with normative age- and sex-specific thresholds for olfactory dysfunction(20). The cost and time taken to administer the full 40-item UPSIT test limits the feasibility of its use in routine clinical and research settings. Hence, several shorter smell identification tests have been developed by Sensonics, either as standalone (two versions of 12-item Brief Smell Identification Test (BSIT)) or preliminary tests (two versions of 4-item Pocket Smell Test (PST)) to guide later administration of the UPSIT to relevant individuals (**see Supplementary Table 1**). A comprehensive list of these and smell tests developed by other companies have been reviewed elsewhere(21).

In this study, we examined the screening performance of Versions A and B of the current 4- item PSTs and 12-item BSITs respectively in a large group of healthy, older individuals from the PREDICT-PD study, and assessed the tests’ ability to detect hyposmia compared with the full 40-item UPSIT. We then sought to identify novel subset(s) of UPSIT items with superior predictive capabilities in the same group, and validate the findings in an independent group of individuals from the same study. We hypothesised that smells from these “winning” subsets could be used as a more accurate pre-screening tool for olfactory dysfunction to guide administration of the full 40-item UPSIT.

## METHODS

### Participant details

We used data from the PREDICT-PD pilot cohort, a study of 1,323 individuals recruited from the general population in the UK between the ages of 60-80. In depth details of recruitment into the PREDICT-PD study has been described elsewhere(22). Of the 1,067 participants from the PREDICT-PD cohort that were sent the full 40-item US version of the UPSIT in the baseline year of the study, 891 completed the test that year (mean age 67.3 years, SD 4.8, 61.5% female). A group of 191 participants who completed the UPSIT test in only Year 3 of the study were used for the validation of “winning” smell subsets (mean age 69.8 years, SD 4.7, 61.8% female). **Figure 1** outlines the workflow of UPSIT data collection from the PREDICT-PD study. A separate group of 44 individuals with established PD (mean age 64.3 years, SD 3.0, 27.3% female) were also sent and completed the full 40-item UPSIT.

**Figure 1:**
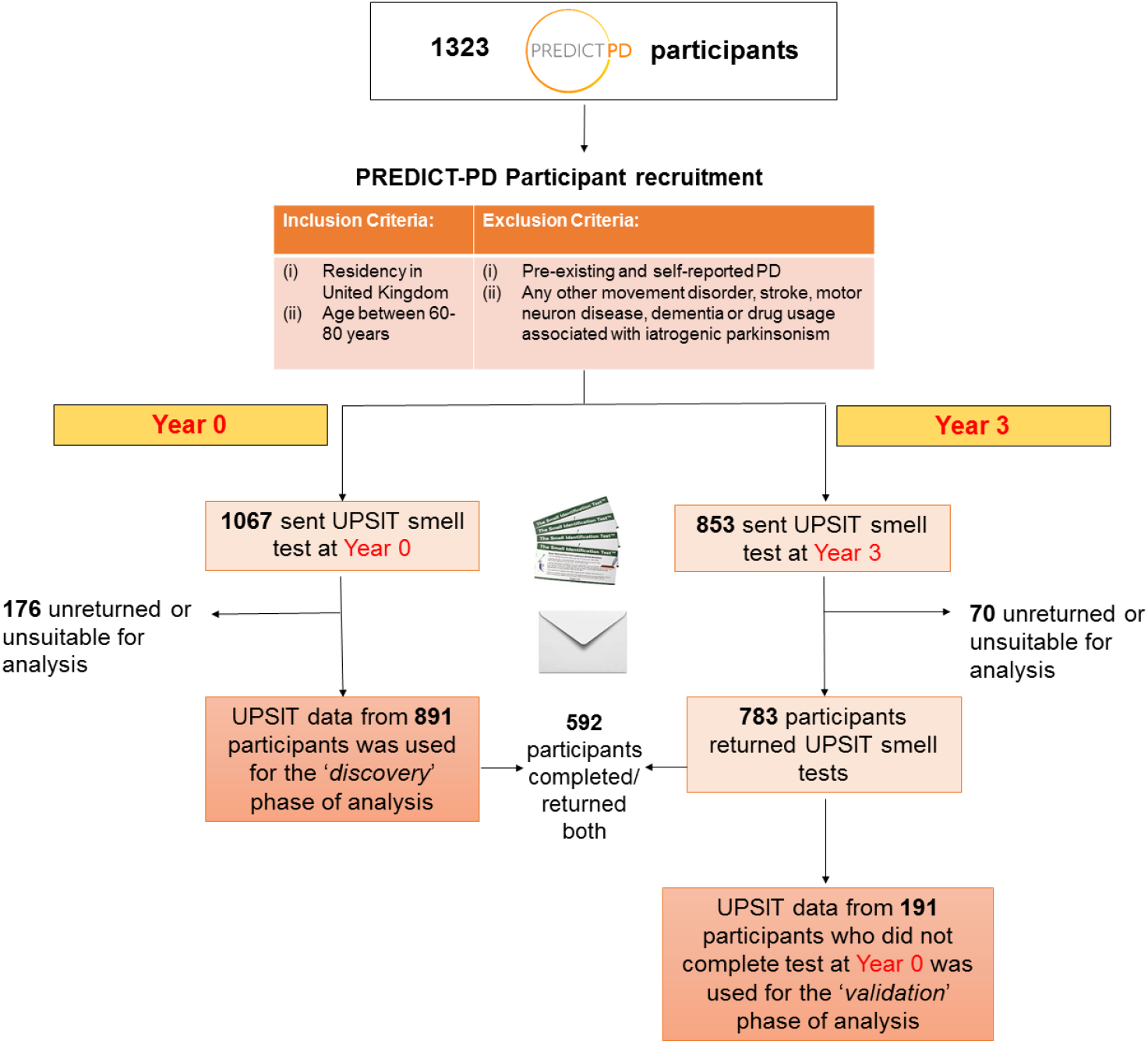
Schematic workflow of PREDICT-PD participation in Year 0 and Year 3 UPSIT smell testing and useable data for ‘*discovery’* and ‘*validation’* cohort smell test analysis

### Assessment of current abbreviated smell tests

Preliminary tests by Sensonics include Versions A and B of the 4-item PST. A test subject is recommended to undergo full UPSIT testing if they cannot correctly identify 1 or more smells in either PST version. The selection of smells in each version was based upon their relevance to diet and nutrition, household, and public safety, rather than empirical evidence relating to smell identification(23). Of the abbreviated standalone tests for olfactory dysfunction by the same company, the BSIT is a validated, cross-cultural 12-item version of the UPSIT(24). In contrast to the PST, the smells within BSIT versions are based upon discriminatory power for specific neurodegenerative diseases: BSIT-A for AD, and BSIT-B for PD.

Scores for all 40 UPSIT smells were recorded for each participant. “True” hyposmia was defined as the lowest 15^th^ centile of UPSIT scores according to gender and age (in 5-year bins), using methods previously described(20). Screening performance of each abbreviated smell test for the detection of hyposmia was assessed against the corresponding total UPSIT scores for each participant. For the 4-item PST and 12-item B-SIT versions, scores of ≤3 and ≤9 respectively are indicative of a positive hyposmia screen. Sensitivity, specificity, positive predictive value (PPV) and negative predictive value (NPV) were calculated for each test.

### Data-driven approach to identify optimal smell subsets

The discovery phase for novel smell item subsets was undertaken using data from the 891 healthy participants (*discovery group*) and assessed all 23,231,378 possible combinations of 1-7 smells from the total of 40 UPSIT smells. For each smell combination, the ability to detect hyposmia was assessed against the full UPSIT score, and was defined in terms of sensitivity, specificity, PPV and NPV, as well as different score thresholds for defining hyposmia. For example, for each of the 18,643,560 combinations of 7 smell subsets from the full set of 40, we assessed screening performance based upon hyposmia being defined as participants scoring 0/7, ≤1/7, ≤2/7, ≤3/7, ≤4/7, ≤5/7 and ≤6/7. The different thresholds for each combination of 7 smell subsets meant we assessed 130,504,920 sets of smell combinations and hyposmia thresholds. Combining this with the same approach for 1-6 smell subsets led to the assessment of a total 157,222,040 possible screening tests.

A “winning” subset of smells at each hyposmia threshold was selected according to those with the highest combined sensitivity and specificity. For example, when considering 5 smell subsets at a threshold of ≤4 to define hyposmia, sensitivity was the number of people who both correctly identified ≤4 of the 5 smells and were defined as hyposmic according to the full UPSIT, divided by the total number of hyposmics according to the full UPSIT. Specificity was the number of people who correctly identified all 5 smells in the subset and were not hyposmic according to the full UPSIT, divided by all those who were not hyposmic as defined by the full UPSIT. These two values were then summed and the combination of smells with the highest combined value was deemed to be the “winner” for that specific threshold. The same process was repeated for every threshold of hyposmia, for all numbers of smell combinations.

Using this method, rather than the area-under the receiver operating curve (AUC), allowed us to identify the best performing combinations of smells across all possible thresholds, rather than one which performed best when averaging across a number of thresholds (as an AUC would). Hence, it allowed us to identify threshold-specific optimal smell subsets and thus enable comparison of different hyposmia thresholds.

The screening performance of each “winning” subset was re-assessed in an independent group of 191 healthy PREDICT-PD participants (*validation group*). There was no overlap in the participants included for selecting the “winning” subsets and the subsequent testing of them (**Figure 1**). Therefore, the results reported are more likely to be generalisable and not due to overfitting of the model.

### Validation of the novel smell subtests in PD individuals

Detection rates of all 40 UPSIT smells were also compared between 44 individuals with PD and the 891 healthy participants. The top discriminating smells were compared with our “winning” smell subsets(s) for assessment of smell overlap. We then compared the screening performance of PST Versions A, B and our “winning” smell subset(s) for hyposmia detection in the 44 individuals with PD.

## RESULTS

Based on total UPSIT scores, 16.2% females (89/548) and 16.0% males (55/343) from the 891 participants in the discovery cohort were classified as having hyposmia. Smoke was the most commonly correctly identified smell (851/891), and Turpentine the least common smell to be correctly identified (328/891). 17.8% and 79.6% of the 191 validation cohort participants and 44 PD participants were classified hyposmic respectively (p<0.001), confirming the recognised greater prevalence of hyposmia in PD patients compared to healthy participants.

### PST and BSIT hyposmia screening performance

**Table 1** displays the screening performances of abbreviated smell tests assessed in the discovery group. Using the recommended cut-off score of ≤3 correctly identified smells to denote hyposmia, PST Version A detected hyposmia with sensitivity 76.8%, specificity 64.9%, PPV 29.3% and NPV 93.6%. PST Version B had a greater sensitivity 88.6% and NPV 94.8%, but lower specificity 45.9% and PPV 23.3%. For the 12-item BSITs, a score of ≤9 on BSIT-A detected hyposmia with a sensitivity of 83.1%, specificity 79.5%, PPV 43.5% and NPV 96.1%. Comparatively, BSIT-B had greater sensitivity 96.5% and NPV 98.7% than BSIT-A, but less specificity 51.8% and PPV 27.5%. We assessed different score thresholds of the BSIT and these are shown in full in **Supplementary Tables 2 and 3**.

**Table 1:** Screening performance of PST and B-SIT Versions A and B for hyposmia detection in discovery cohort.

| Shorter test version | Number of smells | Hyposmia score | Sensitivity | Specificity | PPV | NPV |
| --- | --- | --- | --- | --- | --- | --- |
| PST Version A | 4 | $\leq 3$ | 76.8% | 64.9% | 29.3% | 93.6% |
| PST Version B | 4 | $\leq 3$ | 86.6% | 45.9% | 23.3% | 94.8% |
| BSIT-A | 12 | $\leq 9$ | 83.1% | 79.5% | 43.5% | 96.1% |
| BSIT-B | 12 | $\leq 9$ | 96.5% | 51.8% | 27.5% | 98.7% |

Abbreviations: PPV = Positive predictive value, NPV= Negative Predictive Value

### Identifying optimal smell subsets

We next assessed all combinations of 1-7 smells from the full set of 40 UPSIT smells in the *discovery group*, from which there was a total of 28 “winning” smell combinations. **Table 2** shows a selected set of these “winning” smell combinations and the threshold scores for defining hyposmia. The sensitivity, specificity, PPV and NPV values shown are from their assessment in both discovery and validation groups. The complete results from the data-driven analysis with all 28 “winning” smell combinations at each threshold are presented in full in **Supplementary Tables 4 and 5**, showing their screening performance in the discovery and validation groups separately.

**Table 2:** Screening performance of selected “winning” smell subsets from discovery cohort and re-assessment in validation cohort.

| No. of smells | Hyposmia cut-off score | Smells “winning” in discovery cohort | Sens. in discovery cohort | Spec. in discovery cohort | PPV in discovery cohort | NPV in discovery cohort | Sens.in validation cohort | Spec. in validation cohort | PPV in validation cohort | NPV in validation cohort |
| --- | --- | --- | --- | --- | --- | --- | --- | --- | --- | --- |
| 1 | 0 | Pizza | 69.7 | 66.1 | 28.0 | 92.0 | 82.4 | 53.5 | 27.7 | 93.3 |
| 2 | ≤1 | Clove, Coconut | 62.0 | 88.7 | 50.9 | 92.5 | 61.8 | 82.8 | 43.8 | 90.9 |
| 3 | ≤1 | Pizza, Clove, Root beer | 72.5 | 79.3 | 39.9 | 93.8 | 70.6 | 80.9 | 44.4 | 92.7 |
| 4 | ≤2 | Cherry, Clove, Coconut, Root beer | 72.5 | 87.2 | 51.8 | 94.4 | 61.8 | 88.5 | 53.8 | 91.4 |
| 4 | ≤3 | Menthol, Clove, Gingerbread, Orange | 78.9 | 82.8 | 46.5 | 95.4 | 91.2 | 35.0 | 23.3 | 94.8 |
| 5 | ≤2 | Pizza, Clove, Fruit Punch, Chocolate, Root beer | 71.8 | 88.4 | 54.0 | 94.3 | 73.5 | 84.1 | 50.0 | 93.6 |
| 5 | ≤4 | Menthol, Clove, Onion, Gingerbread, Orange | 83.1 | 80.4 | 44.5 | 96.2 | 94.1 | 33.8 | 23.5 | 96.4 |
| 6 | ≤3 | Pizza, Clove, Fruit Punch, Liquorice, Lime, Pine | 83.8 | 82.6 | 47.8 | 96.4 | 85.3 | 78.3 | 46.0 | 96.1 |
| 6 | ≤4 | Menthol, Cherry, Clove, Gingerbread, Root beer, Orange | 84.5 | 83.8 | 49.8 | 96.6 | 88.2 | 65.6 | 35.7 | 96.3 |
| 7 | ≤4 | Cherry, Mint, Clove, Fruit Punch, Gingerbread, Root beer, Pine | 82.4 | 87.7 | 56.0 | 96.3 | 88.2 | 85.4 | 56.6 | 97.1 |
| 7 | ≤5 | Pizza, Menthol, Cherry, Clove, Gingerbread, Orange, Pine | 87.3 | 83.0 | 49.4 | 97.2 | 100.0 | 53.5 | 31.8 | 100.0 |

*Items in bold include those which most frequently appeared in winning subsets, and in combination*

The “winning” smell subsets have different relative strengths in terms of sensitivity, specificity, PPV and NPV. In both discovery and validation groups, optimised combinations of 7 smells showed superior screening performance to the 12-item BSITs. 7 smell items using a cut-off of 4 for hyposmia surpassed the sensitivity/specificity of BSIT-A (88.2/85.4 vs 83.1/79.5) and 7 smells with a hyposmia cut-off of 5 surpassed that of the more sensitive BSIT-B (100/53.5 vs 96.5/51.8). Using as few as 6 smells could also produce comparable screening performance to the 12-item BSIT (sensitivity/specificity for 6 smells with a cut-off of 3: 85.3/78.3 vs 83.1/79.5 for BSIT-A). Following acquisition of these results, further analysis of smell combinations using >7 UPSIT items was unnecessary and hence were not performed.

For the purpose of selecting a small combination of smells, which would most accurately identify relevant individuals who require further smell testing, it would be important to maximise for sensitivity and NPV in order to minimise the number of impaired individuals missed by further testing. In this regard, just 4 smells (Menthol, Clove, Gingerbread, Orange) with a cut-off of 3 or less produced high sensitivity and NPV scores in both discovery and validation groups when compared with both current 4-item PST tests **(see Table 1, Supplementary Table 4 and Table 2)**. For ease of writing, this particular 4-item subset will henceforth be referred to as version ‘POD’ (PREDICT olfactory dysfunction). Clove featured in almost every “winning” subset of smells, and was the only smell in these “winning” subsets which was already included in the current BSIT or PST tests.

### Comparison of smell identification in PD and healthy participants

Results from comparison of smell identification proportions between PD and healthy participants are shown in **Figure 2**. The four smells featured in version ‘POD’ are among the 7 most discriminating smells, with Menthol and Orange ranked 1^st^ and 2^nd^, with Gingerbread and Clove ranked 5^th^ and 7^th^ respectively.

**Figure 2:**
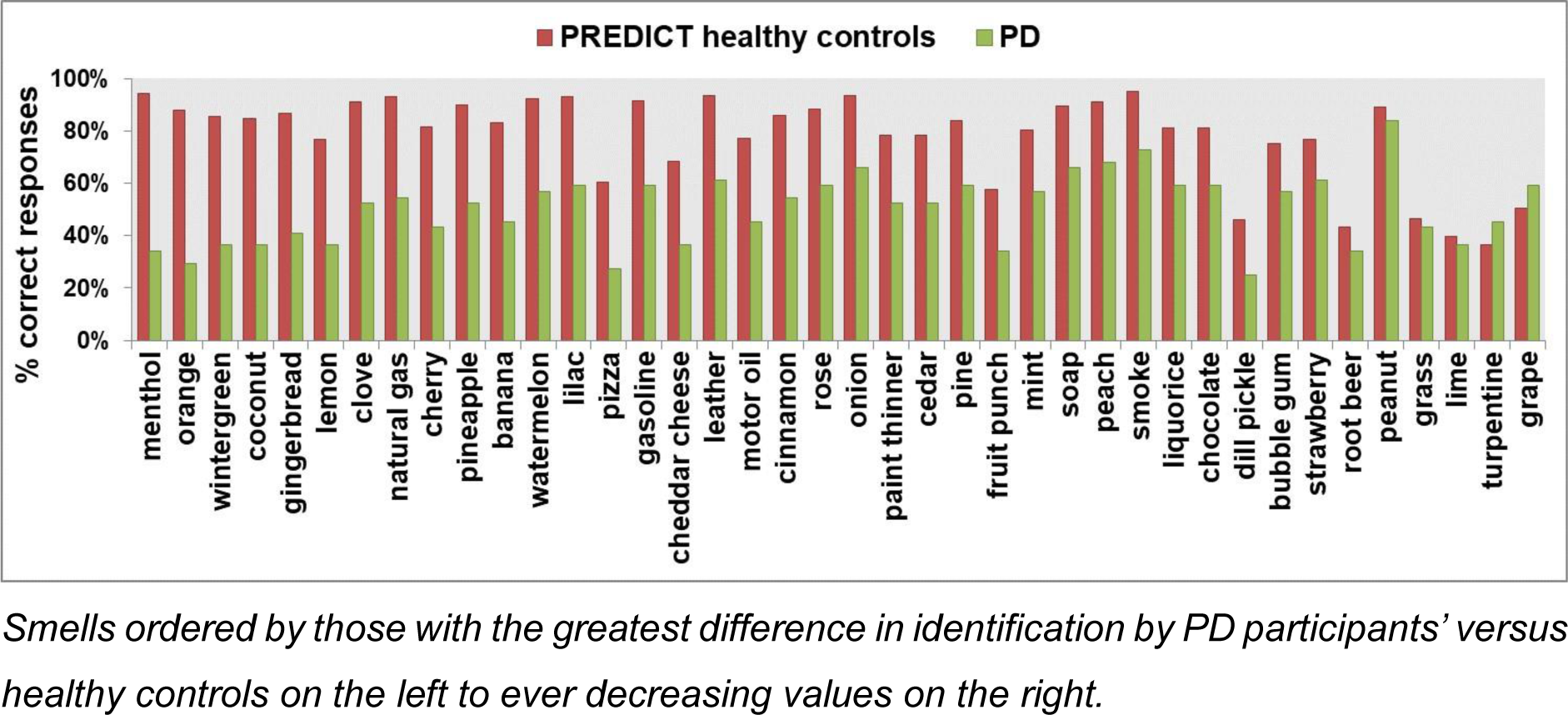
Identification rates of the 40 individual smells in the full UPSIT in 44 PD participants (green) and 891 healthy PREDICT-PD controls (red).

Several smells with lowest discrimination between PD participants and healthy participants (e.g. Grape, Turpentine, Lime and Root beer), generally had the lowest overall smell identification rates in healthy participants. Turpentine and Grape were the only two smells with higher detection by PD participants than healthy participants.

From analysis of 4-item smell tests using ≤3 as the cut-off for olfactory dysfunction in the 44 PD participants, version ‘POD’ identified 100% (35/35) of all PD patients classified with hyposmia based on full UPSIT testing. In contrast, PST Versions A and B respectively identified 85.7% and 74.3% PD patients with hyposmia.

## DISCUSSION

Of the commercially available abbreviated smell identification tests, the 12-item BSITs had predictably greater screening performance for detecting hyposmia than either 4-item PST. This reflects the BSITs’ design to act as standalone shorter tests for hyposmia, whereas PSTs are only intended for use as a pre-screen to target the subsequent administration of full UPSIT testing. The results from our study highlight varying differences in sensitivity and specificity parameters of both 4-item PST Versions, which could impact the overall accuracy of detecting hyposmia, depending on which version is administered.

In our data-driven analysis, we identified subsets of just 7 smells that had superior screening performance compared to the 12-item BSITs when assessed in both discovery and validation phases. As few as 6 smell items produced comparable screening performance to the BSIT, which being half the size of the BSIT could offer obvious benefits in terms of time and expense when undertaking large-scale studies. Previous attempts to devise optimised abbreviated versions of the full UPSIT have produced good screening performance in discovery analyses, but have failed to retain their screening performance when reassessed in independent cohorts(21). The novel smell subsets identified in the present study however, retained their overall screening performance with independent testing.

We also identified a subset of 4 smells (version ‘POD’ - Menthol, Clove, Gingerbread, Orange) which had an overall high screening performance in the discovery cohort, and identified a greater proportion of individuals with hyposmia than either Version A or B of the 4-item PST when reassessed in the independent validation cohort. Whereas, the above 7-item tests may be suitable for standalone testing, this optimised 4-item subset may be an ideal pre-screen test before selective use of the UPSIT.

Studies have consistently demonstrated PD patients to have lower total UPSIT scores compared with healthy controls(19,25),. Here, we were able to interrogate differences between people with PD and healthy controls in greater detail, to look at specific smells which patients with PD might be more likely to lose. All 4 smells included within version ‘POD’ featured among the top 7 discriminating smells between 44 PD and 891 healthy participants, suggesting they may have a particular role in PD-associated olfactory dysfunction. Indeed, when tested in the context of PD, version ‘POD’ was able to correctly identify all 35 PD participants with olfactory dysfunction, outperforming both versions of the PST.

One of the key strengths of the present study is its size. To the best of our knowledge, this is the largest assessment of screening performance of abbreviated versions of smell identification tests in comparison to the full 40-item UPSIT. However, there are a number of limitations. The three ‘distractor’ options used in both PST versions can differ from those for the same smells in the full UPSIT test **(see Supplementary Table 6)**. These different distractors could have influenced participants’ ability to identify the smells to some degree, but the impact is likely to have been relatively small as the target smells are ultimately the same.

Given that the assessment of all of abbreviated smell tests was based upon comparison with participants’ smell status according to their total UPSIT score, we are assuming that it still remains an accurate and sensitive tool for detecting hyposmia in the general healthy population as validated by Doty et al(26). The current study used the US version of the UPSIT, but in a UK population, which might have reduced overall performance due to reduced familiarity with some of the smells. Indeed, a previous UK based study using the US version of UPSIT found certain smells to have low identification rates(27), but this did not include any of the smells from our “winning” subset combinations. The smells with lowest cross-cultural detection included Root beer (52.3%), Lime (56.8%), Dill pickle (61.4%) and Turpentine (65.9%)(27), which was borne out in our own data as both the poorest identified smells in healthy participants as well as the worst discriminating smells between PD participants and healthy participants **(Figure 2).** It is worth noting that these same smells were commonly presented as distractor options for each other’s questions within the full UPSIT test, highlighting the importance of considering the impact of distractor options in influencing individual smell identification scores. Further validation of these findings in larger PD cohorts and in the context of other neurodegenerative diseases such as Alzheimer’s disease would also be useful for future work to explore.

## CONCLUSION

Accurate assessment of olfactory dysfunction may assist in the early detection of certain neurodegenerative diseases such as PD. Using a robust data-driven approach, our study identified several “winning” 1-7 UPSIT smell subsets with high screening performance for hyposmia detection. Of note, 7-item subsets demonstrated superior screening performance to current 12-item BSIT versions. A 4-item subset (Menthol, Orange, Gingerbread and Clove) performed with high sensitivity and NPV for detecting hyposmia both in a general population and specifically in the context of PD; this performance was superior to both current 4-item PST versions and even the standalone 12-item BSIT-A. Significant cost and efficiency savings may be gained by using these smell combinations within an abbreviated smell test to target more focussed administration of the full UPSIT for wider scale clinical and research purposes.

## Acknowledgements

The authors would like to thank and acknowledge all of the participants who have helped support and contribute to the PREDICT-PD project. The authors would also like to remember and acknowledge the valuable contributions from the late Selina Paul to the project, who worked tirelessly to support its aims.

## Contributors

TJ, SA and AJN were involved in the design and conceptualisation of this study, the analysis and interpretation of data, and drafting of the manuscript for intellectual content. LS and DR were responsible for acquisition and cleaning of data, and revising the manuscript for intellectual content. JC, GG, AL, AS were involved in the design and conceptualisation of the PREDICT-PD study and revised the current manuscript for intellectual content.

## REFERENCES

1. Doty RL. Influence of age and age-related diseases on olfactory function. Ann N Y Acad Sci 1989;561:76–86

2. Doty RL. Olfaction in Parkinson’s disease and related disorders. Vol. 46,Neurobiology of Disease 2012;p. 527–52.

3. Haehner A, Boesveldt S, Berendse HW, et al. Prevalence of smell loss in Parkinson’s disease - A multicenter study. Park Relat Disord 2009;15(7):490–4.

4. Nalls MA, McLean CY, Rick J, et al. Diagnosis of Parkinson’s disease on the basis of clinical and genetic classification: a population-based modelling study. Lancet Neurol 2016;14(10):1002–9.

5. Haehner A, Hummel T, Hummel C, et al. Olfactory loss may be a first sign of idiopathic Parkinson’s disease. Mov Disord 2007;22(6):839–42.

6. Chaudhuri KR, Titova N. Nonmotor Parkinson’s: The Hidden Face the Many Hidden Faces Vol. 133, Internation Review of Neurobiology 2017. 1–768.

7. Ross GW, Petrovitch H, Abbott RD, et al. Association of olfactory dysfunction with risk for future Parkinson’s disease. Ann Neurol 2008;63(2):167–73.

8. Berg D, Marek K, Ross GW, et al. Defining at-risk populations for Parkinson’s disease: Lessons from ongoing studies. Mov Disord 2012;27(5):656–65.

9. Siderowf A, Jennings D, Eberly S, et al. Impaired olfaction and other prodromal features in the Parkinson At-Risk Syndrome study. Mov Disord 2012;27(3):406–12.

10. Del Tredici K, Rüb U, De Vos RAI, et al. Where does Parkinson disease pathology begin in the brain. J Neuropathol Exp Neurol 2002;61(5):413–26.

11. Beach TG, Adler CH, Lue LF, et al. Unified staging system for Lewy body disorders: Correlation with nigrostriatal degeneration, cognitive impairment and motor dysfunction. Acta Neuropathol 2009;117(6):613–34.

12. Attems J, Walker L, Jellinger KA. Olfactory bulb involvement in neurodegenerative diseases. Vol. 127, Acta Neuropathologica 2014. p. 459–75.

13. Zou Y, Lu D, Liu L-P, et al. Olfactory dysfunction in Alzheimer’s disease. Neuropsychiatr Dis Treat 2016;12:869–75

14. Roberts RO, Christianson TJH, Kremers WK, et al. Association Between Olfactory Dysfunction and Amnestic Mild Cognitive Impairment and Alzheimer Disease Dementia. JAMA Neurol 2016;73(1):93.

15. Hüttenbrink K-B, Hummel T, Berg D, et al. Olfactory dysfunction: common in later life and early warning of neurodegenerative disease. Dtsch Arztebl Int 2013;110(1–2):1–7, e1.

16. Haehner A, Hummel T, Reichmann H. Olfactory dysfunction as a diagnostic marker for Parkinson’s disease. Vol. 9, Expert Review of Neurotherapeutics 2009. p. 1773–9.

17. Morley JF, Duda JE. Olfaction as a biomarker in Parkinson’s disease. Biomark Med 2010;4(5):661–70.

18. Fullard ME, Morley JF, Duda JE. Olfactory Dysfunction as an Early Biomarker in Parkinson’s Disease. Vol. 33, Neuroscience Bulletin 2017. p. 515–25.

19. Doty RL. Olfactory dysfunction in Parkinson disease. Vol. 8. Nature Reviews Neurology 2012. p. 329–39.

20. Doty RL, Shaman P, Dann M. Development of the university of pennsylvania smell identification test: A standardized microencapsulated test of olfactory function. Physiol Behav 1984;32(3):489–502.

21. Morley JF, Cohen A, Silveira-Moriyama L, et al. Optimizing olfactory testing for the diagnosis of Parkinson’s disease: item analysis of the university of Pennsylvania smell identification test. npj Park Dis 2018;4(1):2.

22. Noyce AJ, Bestwick JP, Silveira-Moriyama L, et al. PREDICT-PD: Identifying risk of Parkinson’s disease in the community: Methods and baseline results. J Neurol Neurosurg Psychiatry 2014;85(1):31–7.

23. Rawal S, Hoffman HJ, Honda M, et al. The Taste and Smell Protocol in the 2011– 2014 US National Health and Nutrition Examination Survey (NHANES): Test–Retest Reliability and Validity Testing. Chemosens Percept 2015;8(3):138–48.

24. Doty RL, Marcus a, Lee WW. Development of the 12-item Cross-Cultural Smell Identification Test (CC-SIT). Laryngoscope 1996;106(3 Pt 1):353–6.

25. Doty RL, Deems D a, Stellar S. Olfactory dysfunction in parkinsonism: a general deficit unrelated to neurologic signs, disease stage, or disease duration. Neurology 1988;38(8):1237–44.

26. Doty RL, Shaman P, Kimmelman CP, et al. University of Pennsylvania Smell Identification Test: a rapid quantitative olfactory function test for the clinic. Laryngoscope 1984;94(2 Pt 1):176–8.

27. Muirhead N, Benjamin E, Saleh H. Is the University of Pennsylvania Smell Identification Test (UPSIT) valid for the UK population? Vol. 6, Otorhinolaryngologist 2013. p. 99–103.

